# Discovery of Membrane Channel Modulators via DNA-Encoded Library Screening Using Native-Like Membrane Protein Nanoparticles

**DOI:** 10.64898/2026.01.27.701919

**Authors:** Francesco V. Reddavide, Trine L. Toft-Bertelsen, Ieva Drulyte, Aspen Rene Gutgsell, Dzung Nguyen, Sara Bonetti, Katerina Vafia, Anne-Sophie Tournillon, Stephan Heiden, Daniel Grosser, Katarina Iric, Veronica Diez, Nanna MacAulay, Stefan Geschwindner, Michael Thompson, Jens Frauenfeld, Robin Löving

**Affiliations:** DyNAbind, Tatzberg 47, 01307 Dresden, Germany; Department of Neuroscience, University of Copenhagen, 2200 Copenhagen, Denmark; Thermo Fisher Scientific, Materials and Structural Analysis, Achtseweg Noord 5, 5651 GG, Eindhoven, The Netherlands. Present address: CryoCloud, Begijnehof 10A, 3512 Utrecht, Netherlands; AstraZeneca, Mechanistic and Structural Biology, Discovery Sciences, R&D, 431 50, Mölndal, Sweden; Salipro Biotech, Teknikringen 38A, 114 28, Stockholm, Sweden; CUP Contract Labs, Carl-Eschebach-Straße 7, 01454 Radeberg, Germany; Institute of Molecular and Cell Biology (IMCB), Agency for Science, Technology, and Research (A*STAR), 61 Biopolis Drive, Proteos, Singapore 138673, Singapore; Nanogami, Koppstraße 16, 81379 München, Germany

## Abstract

Developing novel drugs against membrane proteins is a major challenge in drug discovery due to the difficulty of stabilizing these targets for high-throughput screenings. Pannexin 1 (PANX1) is a membrane channel protein involved in various physiological and pathological processes, making it a promising target for drug discovery. However, efforts to develop PANX1-targeting therapeutics have been hindered by the inherent challenges of stabilizing the protein channel and conducting effective pharmacological screening. Here, we report a proof-of-concept workflow that integrates the Salipro lipid nanoparticle platform with DNA-Encoded Library screenings in a detergent-free format. In this case study, the Salipro DirectMX method was used to generate functional PANX1 nanoparticles for drug discovery and characterisation. Using a high-stringency selection strategy and computational approaches, we identified a specific set of candidate compounds with selective PANX1 enrichment. Surface Plasmon Resonance analysis confirmed the identification of hit compounds. Cryo-Electron Microscopy of the Salipro-PANX1-Compound complex provided structural insights into a potential compound binding site. Electrophysiological recordings in PANX1-expressing *Xenopus laevis* oocytes demonstrated dose-dependent inhibition of PANX1-mediated ion conductance by the compounds. These findings establish a robust workflow for ligand discovery against challenging membrane protein targets and provide novel chemical starting points for the development of PANX1 modulators.

## Introduction

The discovery of novel small-molecule modulators for membrane proteins remains one of the most significant hurdles in drug discovery. Pannexin 1 (PANX1) is a ubiquitously expressed membrane channel protein involved in a wide range of physiological and pathological processes, including ATP release, inflammation, apoptosis, and neuronal signalling^1,2^. Structurally, PANX1 forms large-pore channels that facilitate the passage of ions and small molecules, and its activity is regulated by mechanical stress, post-translational modifications, and interactions with other cellular components^3,4^. Despite its biological importance and therapeutic potential, PANX1 remains a challenging target for drug discovery due to the inherent difficulties formulating stable and functional protein for *in vitro* high-throughput screening.

The DNA-Encoded Library (DEL) technology has emerged as a powerful platform for the discovery of small-molecule ligands against diverse protein targets. DELs enable the screening of millions to billions of compounds in a single experiment by linking each small molecule to a unique DNA barcode^5–7^. However, the application of DEL to membrane proteins has been limited by the need to maintain the native conformation of the purified target throughout the selection process, with only a few successful examples reported to date^8–10^. Traditional detergent-based solubilization often compromises protein stability and function, leading to poor hit quality and high false-positive rates^11^. Cell-based DEL selection has been successfully demonstrated in a few cases^12–15^. These assays avoid the use of detergents, preserving the native cellular environment, but they also introduce significant complexity. The presence of thousands of non-target proteins creates high background noise, which can obscure specific ligand–target interactions and make it difficult to measure binding events accurately and unambiguously.

To address these limitations, we utilized the Salipro DirectMX method, an established membrane protein stabilization technology that enables the direct reconstitution of membrane proteins from cells into Salipro lipid nanoparticles^16–19^. This method preserves a lipid environment that closely resembles native conditions, thereby maintaining the functional integrity of the target membrane protein. Moreover, the incorporation of affinity tags on the saposin scaffold protein enables efficient immobilization of the resulting membrane protein– embedded nanoparticles on solid supports. Altogether, this setup eliminates the need for detergent during downstream screening or analytical procedures^17–19^.

In previous work, we demonstrated the successful structural and functional characterization of Salipro-PANX1 particles using this method^18^. In the present study, we leveraged the technology to perform a DEL screening campaign against PANX1 as a proof-of-concept study with the DyNAbind DEL platform, aiming to identify novel small-molecule binders and functional modulators of this pharmacologically challenging target. Using a combination of high-stringency selection conditions, next-generation sequencing, and computational filtering, we identified a set of candidate compounds with selective PANX1 enrichment. Subsequent validation by surface plasmon resonance (SPR), cryo-electron microscopy (cryo-EM), and electrophysiological assays confirmed the binding and functional inhibition of PANX1 by several hit compounds. These findings demonstrate the feasibility of combining Salipro nanoparticles with DEL screening for membrane protein drug discovery and provide new chemical starting points for the development of PANX1-targeted therapeutics.

## Results

### DNA-Encoded Library Screening Using Salipro-PANX1 Nanoparticles

To identify small-molecule modulators of PANX1, we followed a DEL screening strategy with Salipro-PANX1 nanoparticles schematically illustrated in Figure 1. We transiently expressed a mouse PANX1 (mPANX1) construct in Expi293 cells, which included a cleavable C-terminal GFP and Twin-Strep-tag. The mPANX1 protein was reconstituted using His-tagged Salipro scaffold protein, forming His-Salipro-mPANX1 nanoparticles. These were purified via Strep-Tactin Sepharose affinity chromatography, with on-column cleavage by PreScission protease, following previously established protocols^18^. Biochemical characterization of the resulting His-Salipro-mPANX1 particles is shown in Supplementary Figure 1A. The DEL campaign utilized DyNAbind libraries comprising over 11 million compounds synthesized via two-, and three-cycle split-and-pool amide bond chemistry^20^. Selections were performed under three separate conditions: (i) no-protein control (NPC), (ii) lipid-only His-Salipro nanoparticles (His-Salipro-Empty) control, and (iii) His-Salipro-mPANX1 in the presence of a 10-fold molar excess of untagged Salipro-Empty particles (His-Salipro-mPANX + Salipro-Empty). The final condition was designed to increase selection stringency by outcompeting non-specific nanoparticle binders while retaining PANX1-specific interactions.

**Figure 1.**
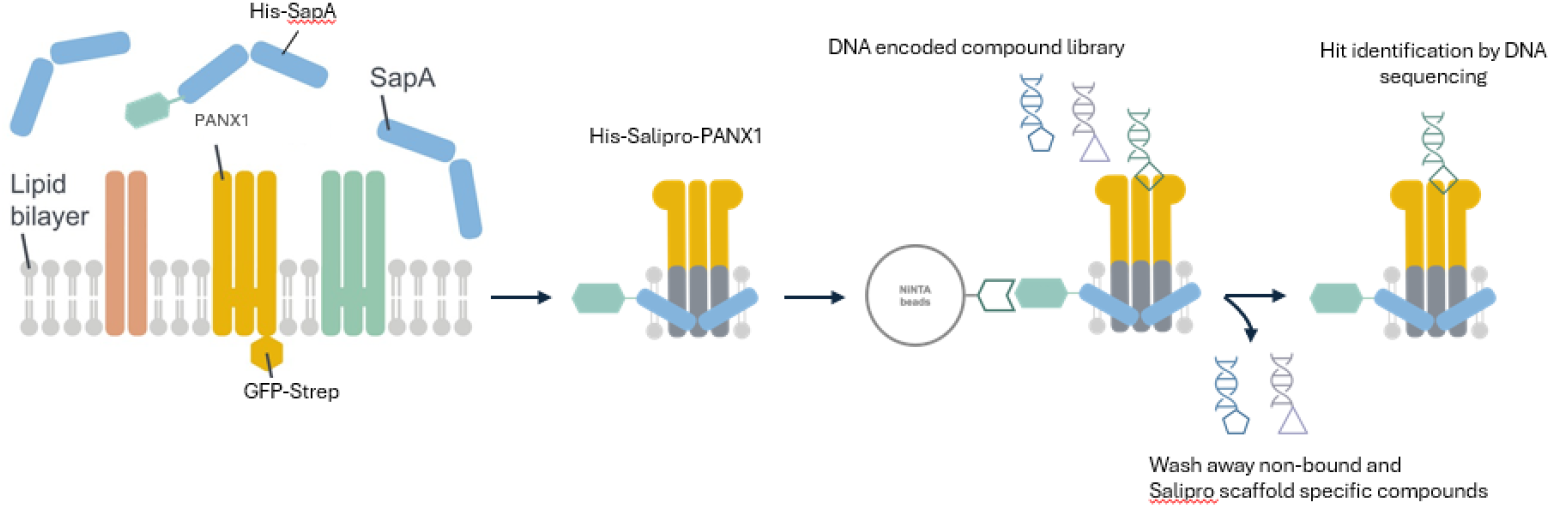
Schematic overview of the Salipro-DirectMX and DEL workflow. His-Salipro-mPANX1 nanoparticles were assembled from a mixture of His-tagged and non-tagged saposin scaffold proteins and purified via cleavable GFP with a C-terminal Twin-Strep-Tag. For DEL selection, particles were immobilized on Ni-NTA magnetic beads, and a 10-fold excess of non-tagged Salipro-Empty particles was added to absorb scaffold binders, which were removed along with non-bound compounds during washing. After iterative rounds, mPANX1-enriched binders were PCR-amplified and identified by next-generation sequencing.

His-tag pull down magnetic beads were used to immobilize His-Salipro particles, allowing detergent micelle-free DEL screening of the wildtype and tag-free mPANX1 channel protein. Following iterative rounds of selection, eluates were amplified by PCR and analyzed via next-generation sequencing (NGS). The sequencing data were processed using a custom KNIME-based pipeline, and enrichment scores were normalized across library sizes and experimental conditions^21^. Compounds exhibiting strong enrichment in the His-Salipro-mPANX1 + Salipro-Empty condition (iii), with minimal enrichment in the NPC (i) and His-Salipro-Empty (ii) control conditions.

From the enriched pool, 5,853 compounds were virtually synthesized and analysed for physicochemical properties and structural fingerprints. Structural clustering was performed using Morgan fingerprints and a Tanimoto similarity threshold of 0.8, resulting in chemically related clusters. Filtering based on enrichment profiles and structural redundancy yielded a refined set of 1,084 candidate compounds.

To accelerate hit validation, a similarity screen was conducted against a 2-million-compound drug-like library from Mcule. identifying 97 structurally similar, commercially available compounds by using substructure fingerprints, FragFP in Datawarrior^22,23^. The DEL selection workflow is schematically outlined in Supplementary Figure 2. Molecular docking against both mouse (PDB 8A3B) and human (PDB 6LTO) PANX1 structures suggested preferential binding to the tryptophan belt and inner pore region (Supplementary Figure 3) indicating binding to a functionally important sites involved in channel gating and inhibitor sensitivity^24–27^. From these hits, nine compounds with favourable docking scores were selected for experimental validation.

### SPR and Cryo-EM Verify PANX1 Binding Hits

To confirm binding of the selected compounds to PANX1, we performed surface plasmon resonance (SPR) analysis. His-Salipro-mPANX1 particles were immobilized on Ni-NTA sensor chips, with His-Salipro-Empty particles serving as a background reference. Of the nine compounds tested, two exhibited reproducible binding with dissociation constants (Kd) in the micromolar range: Compound 5 (estimated Kd 190 µM; Figure 2A), and Compound 6 (estimated Kd 102 µM; Figure 2B). Compound 2 also appeared to bind mPANX1 but had complex binding behaviour on the sensor surface that prevented confident KD determination (data not shown). We further evaluated compound binding to human PANX1 (hPANX1), reconstituted into Biotin-Salipro-hPANX1 particles (Supplementary Figure 1B). Compound 5 (Kd 53 µM) and Compound 6 (Kd 43 µM) exhibited affinities comparable to those observed with mPANX1 (Supplementary Figure 4A-B). As a control experiment, measurements with the known ligand Benzoylbenzoyl-ATP (BzATP) were performed. These displayed weak and transient binding to both His-Salipro-mPANX1 (Kd 5.0 mM; Figure 2C) and Biotin-Salipro-hPANX1 (Kd 1.3 mM; Supplementary Figure 4C), consistent with previous reports validating the SPR setup^18^.

**Figure 2.**
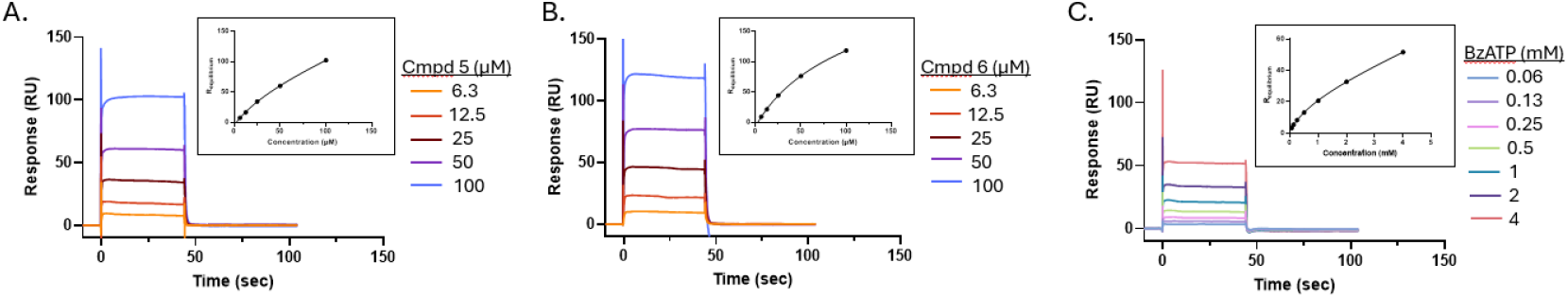
Hit Compound Binding Profiles for mPANX1. Concentration series of (A) compound 5, (B) compound 6, and (C) benzoylbenzoyl-ATP were injected over a mPANX1 coated surface for 45 seconds followed by a 60 second dissociation time. Equilibrium analysis for each dataset is shown as an inset. Data is representative of n=2.

To visualize the binding mode of Compound 6, we performed single-particle cryo-Electron Microscopy (cryo-EM) on the Salipro-mPANX1-Compound 6 complex, yielding a resolution of 4.5 Å, while no symmetry averaging was applied during reconstruction (Fig. 3A). As observed in the apo structure, the extracellular and transmembrane regions were better resolved than the intracellular domain, likely due to intrinsic flexibility (Supplementary Figure 5). Rigid-body docking of the apo PANX1 atomic coordinates (PDB: 8A3B) revealed minimal conformational changes upon compound binding. A putative electron density was observed within the hydrophobic pore, likely corresponding to Compound 6 in close proximity to Ser73 and Trp74 (Fig. 3B). Trp74 has previously been associated with channel gating, suggesting a potential functional relevance of this binding site^24–27^. However, given the modest resolution of the map, no further atomic coordinate refinement or structural interpretation was performed.

**Figure 3.**
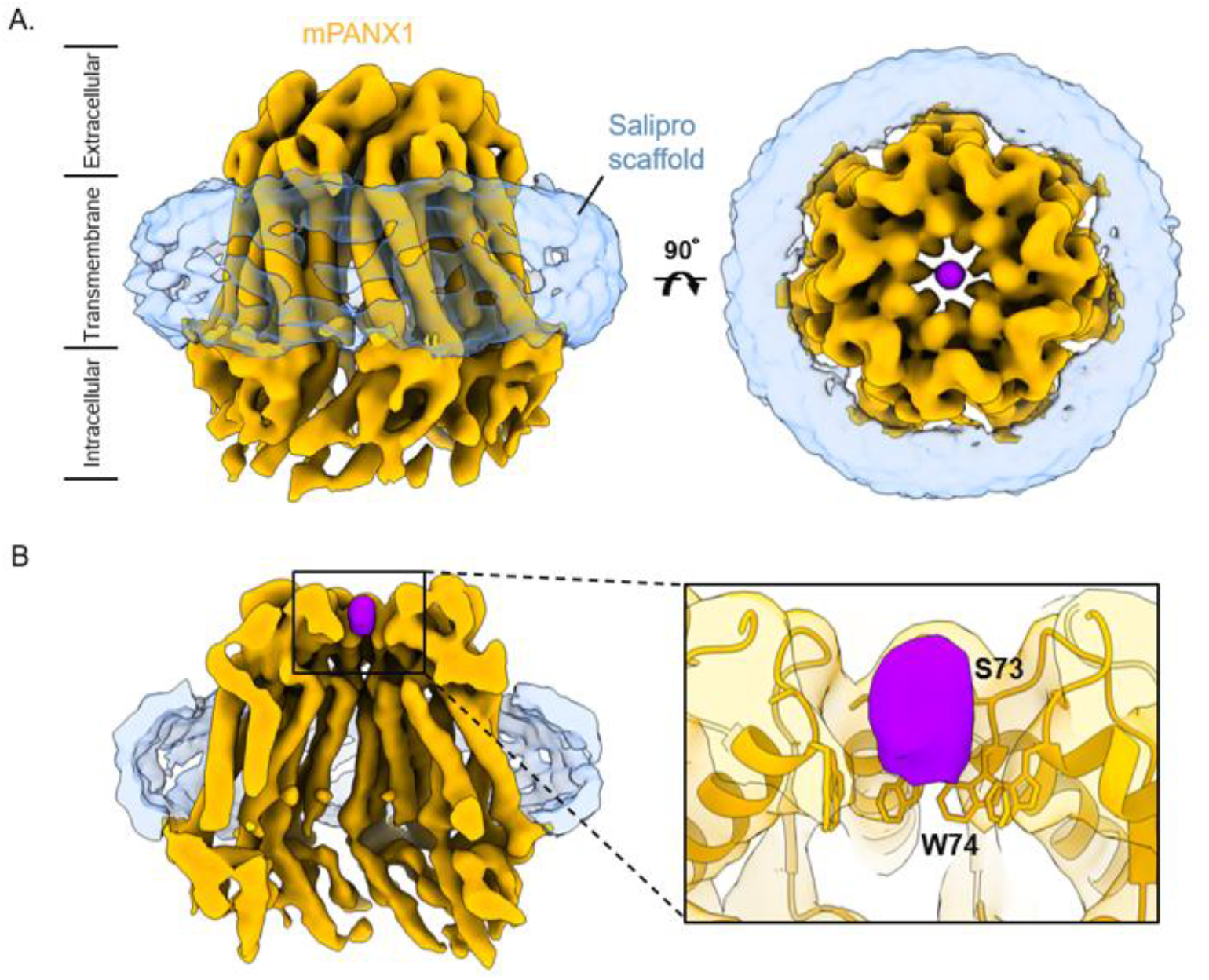
Cryo-EM analysis of Salipro-mPANX1-Compound 6. (A) Unsharpened cryo-EM map of Salipro-mPANX1-Compound 6 reconstruction at 4.5 Å resolution. The map was calculated without imposing any symmetry averaging. The ion channel (orange) and Salipro lipid disk (blue) are shown as side view (left) and top view (right). The compound density is shown in purple. (B) Central slide through the reconstruction showing the compound density in the ion channel pore. The inset displays the zoomed-in region with the residues closest to the compound labelled. The ion channel and compound densities are shown at the same threshold level.

### Functional Characterization of PANX1 Inhibition

To assess the functional impact of the identified compounds, we heterologously expressed rat PANX1 (rPANX1) in *Xenopus laevis* oocytes and performed two-electrode voltage clamp recordings as previously described^28^. rPANX1-expressing oocytes exhibited robust outwardly rectifying ion currents compared to uninjected control oocytes (P < 0.001, Fig. 4A). The rPANX1-mediated current was inhibited by the well-established PANX1 inhibitor carbenoxolone^28^ (P < 0.01, n = 6, Figure 4A) verifying successful rPANX1 expression in the oocytes.

**Figure 4.**
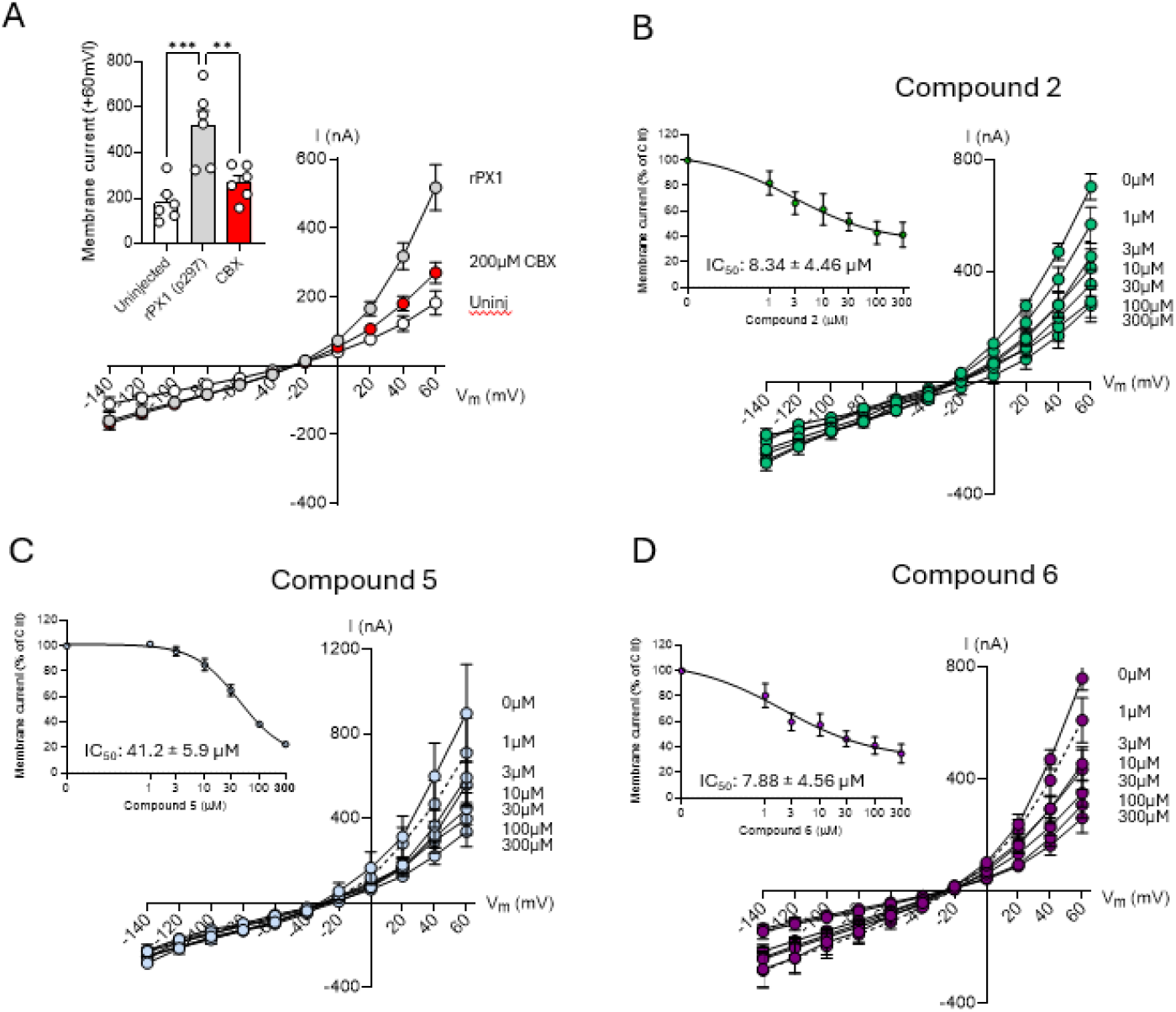
Functional characterization of PANX1 hit compounds. (A) Summarized I–V relationships from uninjected oocytes, Panx1-expressing oocytes and Panx1-expressing oocytes exposed to 200 µM carbenoloxone (CBX). Inset: Summarized and averaged membrane current at +60 mV measured. B-D, summarized I–V relationships from Panx1-expressing oocytes (n = 5-6) treated with compound 2 (B), compound 5 (C) or compound 6 (D) during the voltage step protocol. Membrane currents measured at +60 mV were used for the dose–response curve analysis (B-D, insets). Data were fitted with GraphPad Prism using non-linear regression analysis (B-D, insets). To test for statistical significance, a one-way ANOVA was used (A). **P < 0.01; ***P < 0.001. All data points are represented as means ± SEM.

We next evaluated the effect of Compounds 2, 5, and 6 on rPANX1-mediated ion conductance. All three compounds inhibited rPANX1 currents in a dose-dependent manner (Fig. 4B-D), with IC_50_s of 8.34 ± 4.46 µM for compound 2, 41.2 ± 5.90 µM for compound 5, and 7.88 ± 4.56 µM for compound 6, n = 5-6, Fig. 4B-D, insets). These results confirm that the identified compounds not only bind rPANX1 but also modulate its ion channel function.

## Discussion

The identification of small-molecule modulators for membrane proteins such as PANX1 remains a significant challenge in drug discovery, largely due to the inherent instability of these targets outside their native lipid environment^19,29,30^. In this study, we demonstrate the successful integration of the Salipro-DirectMX platform with with the DyNAbind DEL platform to overcome these limitations and identify novel PANX1-binding compounds with functional activity. The Salipro-DirectMX approach enabled the direct reconstitution of PANX1 channel from cells into lipid nanoparticles, preserving its native conformation and functional integrity. The detergent-free DEL setup was critical for maintaining the native structural and functional properties of PANX1 throughout the selection process. The use of affinity tags on the Salipro scaffold allowed for efficient immobilization of unmodified PANX1 on solid supports, facilitating both DEL screening and downstream biophysical assays such as SPR.

The DEL campaign employed a high-stringency selection strategy, incorporating multiple controls and iterative enrichment, to reduce off-target binding and enhance hit specificity. Using immobilized His-Salipro-mPANX1 particles alongside a tenfold molar excess of untagged Salipro-Empty particles effectively enriched for PANX1-specific binders. Subsequent integration of Next Generation Sequencing, structural clustering, and virtual screening enabled prioritization of a focused set of candidate compounds. Notably, off-the-shelf compound matching and molecular docking streamlined the transition from hit identification to experimental validation, eliminating the need for immediate hit resynthesis. SPR analysis confirmed that three compounds bind to Salipro-mPANX1 particles, with compounds 5 and 6 showing verified dissociation constants in the low-to mid-micromolar range.

These affinities are typical for initial hit-stage ligands and are in the same range as those reported for known PANX1 antagonists, such as spironolactone (Kd 160 µM) and carbenoxolone (Kd > 256 µM)^18^, providing a promising starting point for further optimization.

Cryo-EM analysis of the Salipro-mPANX1–Compound 6 complex revealed a putative electron density within the ion channel pore, localized near residues Ser73 and Trp74 of PANX1 (Figure 3B). This region has previously been implicated in channel gating and inhibitor interaction^24–27^. The potential density of Compound 6 in this region supports its role as a modulator of PANX1 activity and underscores the pharmacological relevance of the extracellular loop interface. Due to the modest resolution, extensive atomic refinement was not pursued. However, further analysis using orthogonal validation techniques are part of future studies to confirm the binding site and mechanism.

The expression of rPANX1 in *Xenopus laevis* oocytes has previously been used for PANX1 functional characterization and determination of inhibitory action of known PANX1 inhibitors^28^. Our electrophysiological recordings in the oocytes confirmed that the three hit compounds inhibited rPANX1-mediated ion conductance in a dose-dependent manner with IC_50_s in the 10-40 µM range. These findings validate the functional relevance of the identified hits and support their potential as starting points for the development of PANX1-targeted therapeutics. Future work will focus on SAR studies, orthogonal validation of binding sites, and in vivo evaluation of the identified compounds to further explore their therapeutic potential.

Taken together, our results highlight the utility of combining Salipro-DirectMX with DEL screening for the discovery of ligands targeting challenging membrane proteins. This integrated approach enables the identification of functionally active compounds while preserving the structural integrity of the target, addressing a key bottleneck in membrane protein drug discovery.

## Methods

### Material and reagents

Human saposin A was recombinantly produced and purified as previously described^16,31^. Pure saposin A in HN buffer (50 mM HEPES, pH 7.5, 150 mM NaCl,) was stored at −80 °C. Saposin A empty nanoparticles (Salipro-Empty particles) were prepared as previously described^16,18^. To generate affinity-tagged Salipro-Empty particles, a mixture of scaffold protein was used in which one-third carried the affinity tag and two-thirds were tag-free, aiming to achieve approximately one tag per reconstituted particle. PreScission protease was purchased from Cytiva. PANX1 ligands for SPR binding measurements were Bz-ATP (Sigma-Aldrich, CAS 112898-15-4) and Carbenoxolone (sourced at AstraZeneca).

### PANX1 Expression

Mouse PANX1 (mPANX1) and human PANX1 (hPANX1) constructs have been described previously^18^. Briefly, mPANX1 was cloned into the pcDNA3.4 TOPO™ vector with an N-terminal FLAG-tag and a C-terminal GFP fused to a Twin-Strep-tag, flanked by PreScission protease cleavage sites (FLAG-3C-mPANX1-3C-GFP-TwinStrep), allowing upon PreScission (Cytiva) cleavage, isolation of tag-free mPANX1. The construct was codon-optimized for expression in human Expi293 cells, which were transiently transfected and harvested five days post-transfection. The hPANX1 construct was cloned into the pFastBac1 vector with a C-terminal eGFP, His_10_-tag, and EPEA affinity tag, along with a PreScission cleavage site (hPANX1-3C-GFP-His_10_-EPEA), enabling isolation of tag-free hPANX1 after PreSission cleavage. Codon-optimized for insect cells, hPANX1 was expressed in Sf9 cells following baculovirus infection, with cell pellets collected 48 h post-infection.

For *Xenopus* laevis oocyte expression, cDNA encoding rat PANX1 (rPANX1), originally provided by R. Bruzzone^32^ and cloned into the pXOOM vector^33^, was linearized and transcribed into cRNA using the T7 mMessage Machine kit (Thermo Fisher Scientific). The cRNA was purified with MEGAclear and stored at −80 °C. Oocytes were maintained in Kulori medium (90 mM NaCl, 1 mM KCl, 1 mM MgCl_2_, 1 mM CaCl_2_, 5 mM HEPES, pH 7.4 adjusted with 2 M Tris-base) at 19 °C for 24 h prior to injection. Each oocyte was microinjected with 13 ng of rPANX1 cRNA and incubated for 3–4 days in Kulori medium at 19 °C before electrophysiological experiments^28^.

### Purification of Salipro-PANX1 Particles

Salipro-mPANX1 and Salipro-hPANX1 particles were purified and characterized as previously described^18^. Briefly, for Salipro-mPANX1, 500 µL of Strep-Tactin Sepharose High Performance resin (GE Healthcare) were equilibrated with HNG buffer in a Poly-prep® chromatography column (Bio-Rad). Lysate from a 2.5 g Expi293 cell pellet was loaded onto the column, and the flow-through was passed three additional times over the resin. The PANX1-bound resin was incubated with 45 mL HNG buffer containing 1 mg/mL saposin A for 30 min at 4 °C. After discarding the flow-through, the resin was washed with 16 column volumes (CV) of HNG buffer. Bound Salipro-PANX1-GFP was cleaved overnight at 4 °C using 60 U of PreScission protease (Cytiva) in 600 µL HNG buffer. The cleaved sample was reloaded into the column, and tag-free Salipro-mPANX1 particles were eluted in the flow-through using 3 CV HNG buffer. The eluted particles were concentrated using a 50-kDa NMWL Amicon filter (Millipore) and subjected to preparative size-exclusion chromatography (SEC) on a Superose 6 Increase 10/300 GL column (Cytiva), equilibrated in HNG buffer, using an Äkta Pure system at a flow rate of 0.35 mL/min. Fractions were analyzed by SDS-PAGE (NuPAGE 4–12% Bis–Tris gels, Thermo Fisher Scientific) and visualized by silver staining. Pure fractions were pooled, concentrated, flash-frozen, and stored at −80 °C.

For Salipro-hPANX1, lysate from 1 g Sf9 cell pellet was loaded onto a 300 µL CaptureSelect™ C-tag Affinity Matrix (Thermo Fisher Scientific), and the flow-through was passed three additional times over the resin. The resin was incubated with 24 mL HNG buffer containing 1 mg/mL saposin A for 30 min at 4 °C, followed by washing with 20 CV HNG buffer. Bound Salipro-hPANX1 was cleaved overnight at 4 °C using 60 U of PreScission protease in 400 µL HNG buffer. The cleaved sample was reloaded into the column, and tag-free Salipro-hPANX1 particles were eluted in the flow-through using 3 CV HNG buffer. Concentration, SEC purification, and fraction analysis were performed as described for Salipro-mPANX1.

### DNA-Encoded Library (DEL) Selection

To identify small-molecule binders to PANX1, a DEL selection was performed using His-Salipro-mPANX1 nanoparticles. The combined DEL, generated via two- or three-cycle split-and-pool chemistry, contained over 11 million unique compounds. Selections were carried out on His-tag magnetic beads using a KingFisher magnetic bead handler. Iterative selection rounds were performed, using eluates from each round as input for the next. After the final round, bound DNA tags were PCR-amplified and sequenced by Illumina NGS, with data processed through a custom KNIME pipeline^34,35^. Reads were demultiplexed according to the unique index assigned to each selection, and the code region was extracted and mapped to the corresponding identity. Code frequencies per sample were determined by counting the occurrences of each code across all reads of each sample. Enrichment was calculated by normalizing code counts to total read counts to obtain relative abundances, followed by dividing post-selection abundance by pre-selection abundance to assess distributional changes.

Decoding and deconvolution were carried out based on the correct index unique to each selection, with correction for 1–2 mutation errors. The coding region was then extracted from each sequence, incorporating the correction of single-base mutations, and assigned to its corresponding identity. The number of reads corresponding to each specific coding region and experimental condition was quantified. The normalized Zn score for each compound was calculated using the formula developed by Faver et al.^36^. A Zn score above 0 indicates that the structure is enriched beyond what would be expected by random chance, provided that enrichment does not occur under EP and NP conditions. A structure with a Zn score above 0 in the target condition and below 0 in the non-target condition is considered enriched. Compounds showing strong enrichment in condition (iii) His-Salipro-mPANX1 + Salipro-Empty condition with minimal signal in control conditions were prioritized as potential binders.

The most highly enriched structures were selected for further examination as potential binders. To improve statistical robustness, we simulated oversampling by grouping detections for substructures. For example, in a two-cycle library, we aggregated counts for individual substructures across cycle 1 and cycle 2. The hits with the highest enrichment were then fingerprinted and mapped onto clusters of chemically related compounds (using Morgan fingerprints and a Tanimoto similarity threshold of 0.8), also an off-the-shelf 2-million-compound drug-like library from Mcule (www.mcule.com) was screened for similarity to facilitate final hit selection and further validation analysis. The visualization of the similarity plot was done in DataWarrior software based on the substructure fragment dictionary-based fingerprint descriptor FragFp.

The enriched compounds were subjected to molecular docking. Nine compounds with favourable docking scores were selected for purchase from Mcule and subsequent experimental validation.

### Surface Plasmon Resonance (SPR) Details

All SPR experiments were conducted on a BIAcore S200 or T200 instrument (Cytiva) using 10 mM HEPES, pH 7.4, 150 mM NaCl (HBS-N) as the running buffer.

#### His-Salipro-mPANX1 SPR Assays

Multicycle kinetic experiments were conducted using a sensor with Ni^2+^ ions complexed on a two-dimensional chelating surface (NiHC 1500M, Xantec Bioanalytics GmbH) at 12 °C. Prior to ligand immobilization, the sensor was washed with 300 mM EDTA, pH 8.3, and loaded with Ni^2+^-ions by injecting a 500 nM solution of NiCl_2_ in running buffer for 3 min. His-Salipro-mPANX1 and His-Salipro-Empty particles were injected at a concentration of 10–20 ng/mL using a contact time of 16 min at 1 µL/min to achieve desired densities of 2500–4000 RU. His-Salipro-Empty particles were immobilized on the reference surface to serve as a control for non-specific binding. Increasing concentrations of (3.1, 6.3, 12.5, 25, 50, and 100 µM) of hit compounds or (0.13, 0.25, 0.50, 1.00, 2.00, 4.00 mM) of BzATP were iteratively injected with a 45 s contact time, followed by a 60 s dissociation phase. All resulting sensorgrams were reference and blank subtracted prior to fitting.

#### Biotin-Salipro-hPANX1 SPR Assays

Multicycle kinetic experiments were conducted using a streptavidin-coated sensor (Cytiva) at 12 °C. Prior to ligand immobilization, the sensor was washed with 50 mM NaOH, 1 M NaCl for 1 min. Biotin-Salipro-hPANX1 and Biotin-Salipro-Empty particles were injected at a concentration of 10–20 ng/mL using a contact time of 16 min at 1 µL/min to achieve desired densities of 2500–4000 RU. Biotin-Salipro-Empty particles were immobilized on the reference surface to serve as a control for non-specific binding. Increasing concentrations of (3.1, 6.3, 12.5, 25, 50, and 100 µM) of hit compounds or (0.13, 0.25, 0.50, 1.00, 2.00, 4.00 mM) of Bz-ATP were iteratively injected with a 60 s contact time, followed by a 90 s dissociation phase. All resulting sensorgrams were reference and blank subtracted prior to fitting.

### Cryo-EM sample preparation and data collection

Quantifoil® R 1.2/1.3 Ultrathin Carbon grids were glow-discharged using 5 mA current for 30 s (GloQube®, Quorum Technologies). Compound 6 was added to purified Salipro-mPANX1 particles at a final concentration of 500 μM. To overcome preferred orientation, protein sample was mixed with 0.5 mM Fos-Choline 8 (Anatrace) before the grid freezing. 3 μL of protein:ligand sample were pipetted onto the grid in Vitrobot Mark IV (Thermo Fisher Scientific) chamber set to 4 °C and 95% humidity. Grids were then blotted for 5 s using a blot force of 0 and 30 s waiting time before plunge-freezing in liquid ethane.

Data collection was conducted using a 200 kV Thermo Scientific™ Glacios™ 2 Cryo-Transmission Electron Microscope (Cryo-TEM) equipped with Selectris Imaging Filter and Falcon 4i Direct Electron Detector camera operated in Electron-Event representation (EER) mode. Thermo Fisher Scientific EPU 3 software was used to automate the data collection. Aberration-free image shifts (AFIS) and Fringe-free Imaging (FFI) increased the data collection throughput to ∼520 movies/hour. 30 µm C2 aperture was utilised to limit the parallel beam diameter to ∼1 µm. 10,000-movie dataset was collected in EFTEM mode using 10 eV slit and a nominal magnification of 165,000×, resulting in a calibrated pixel size of 0.694 Å. The exposure time of 2.29 s with a total dose of 49.8 e^-^/Å^2^ was used, and each movie was split into 50 fractions during motion correction. The dose rate on the camera was 10.8 e^-^/px/s, and the nominal defocus range was specified between −0.75 and −1.5 µm in 0.25 µm intervals. Data collection parameters are listed in Supplementary Table 1.

### Cryo-EM data processing and structural analysis

Data processing was performed using Relion^37^ and CryoSPARC^38^, as described previously^18^. In brief, after motion- and CTF-correction (using Relion’s implementation of MOTIONCOR^39^ and CTFFIND-4.1^40^, respectively), particles were picked using a template-free auto-picking procedure based on a Laplacian-of-Gaussian (LoG) filter with particle diameter set to 130–170 Å. A total of 1,367,517 particles from 9299 micrographs were extracted in a 320 px box Fourier cropped to 160 px box. Following 2D and 3D classification, the best 3D class, consisting of 327,491 particles, was chosen. Refinement using C1 symmetry, and masking and sharpening resulted in the final 4.5 Å reconstruction. The map resolution was determined based on the gold-standard 0.143 criterion.

Atomic coordinates for the previously published Salipro-mPANX1 structure (8A3B.pdb) were rigid-body fitted into the density of the Compound 6 bound structure using ChimeraX^41^. Local resolution was estimated using Relion.

### Electrophysiology measurements

Conventional two-electrode voltage clamp studies were performed using a DAGAN CA-1B High-Performance oocyte clamp (DAGAN, Minneapolis, MN, USA) with a Digidata 1440A interface controlled by pCLAMP software, version 10.5 (Molecular Devices, Burlingame, CA, USA). Electrodes were pulled (HEKA Elektronik, Lambrecht, Germany) from borosilicate glass capillaries to a resistance of 2.5–4 MΩ when filled with 1 M KCl. The currents were low pass filtered at 500 Hz and sampled at 1 kHz. Current measurements were derived from an 11-step voltage clamp protocol (200 ms, 20 mV increments from −140 to +60 mV) with a holding potential of −30 mV. The recording solution contained: 100 mM NaCl, 2 mM KCl, 1 mM CaCl_2_, 1 mM MgCl_2_, 10 mM HEPES, adjusted to a pH of 7.4 with 2 M Tris-base. Stock solutions of carbenoxolone (50 mM), Compound 2, Compound 5, and Compound 6 (5 mM) were all prepared in ddH_2_O and employed for the subsequent drug dilutions (with same total amount of vehicle in all employed concentration). The oocytes were exposed to a given compound concentration for 5 min prior to recording. Oocyte experiments were from two different frog donors. The statistical analysis (one-way ANOVA) was performed with Prism 9 (GraphPad Software Inc., La Jolla, CA, USA). Membrane currents measured at +60 mV were used for dose–response curve analysis. Data were fitted with GraphPad Prism using non-linear regression analysis to obtain IC_50_ concentrations of compounds. Data represent the means ± SEM and P<0.05 was considered statistically significant.

## Data availability

The cryo-EM density map has been deposited in the Electron Microscopy Data Bank with the accession code EMD-56448.

