## Supplementary figures for "Discovery of Membrane Channel Modulators via DNA-Encoded Library Screening Using Native-Like Membrane Protein Nanoparticles"

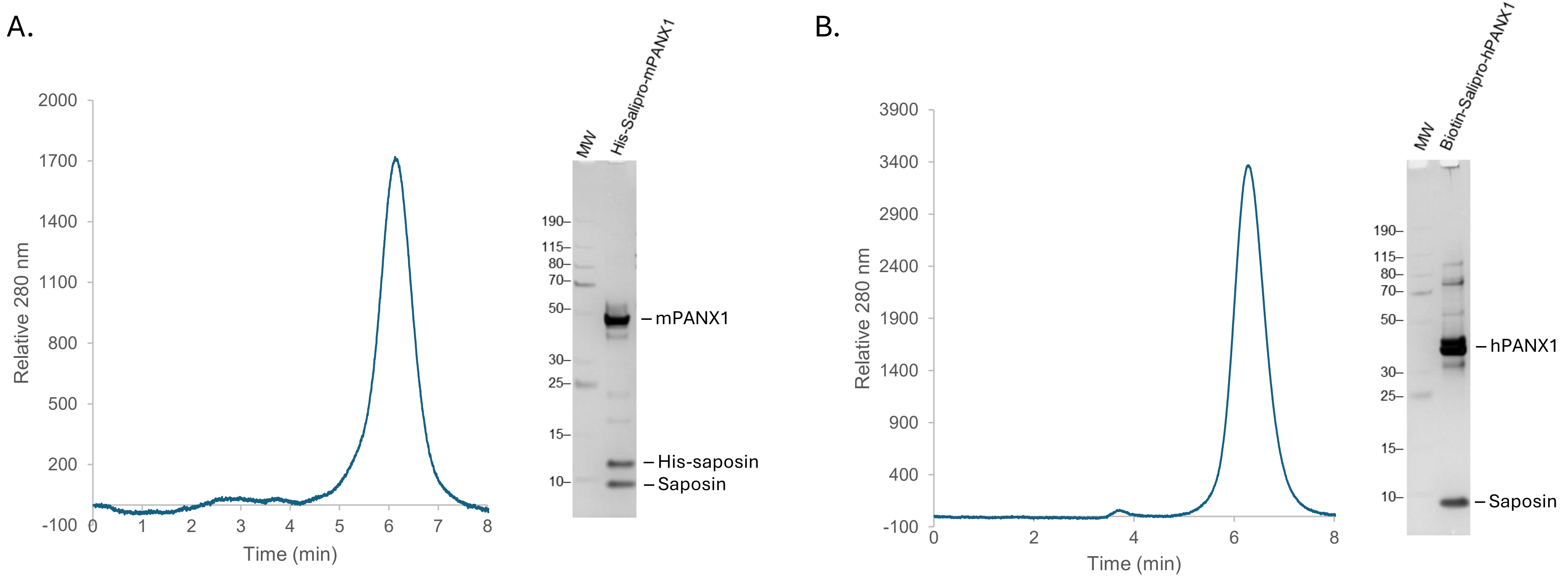


**Supplementary Figure 1.** Salipro-PANX1 nanoparticle production. Analytic SEC and SDS-PAGE of purified (A) His-Salipro-mPANX1 and (B) biotin-Salipro-hPANX1 confirms nanoparticle purity and homogeneity.

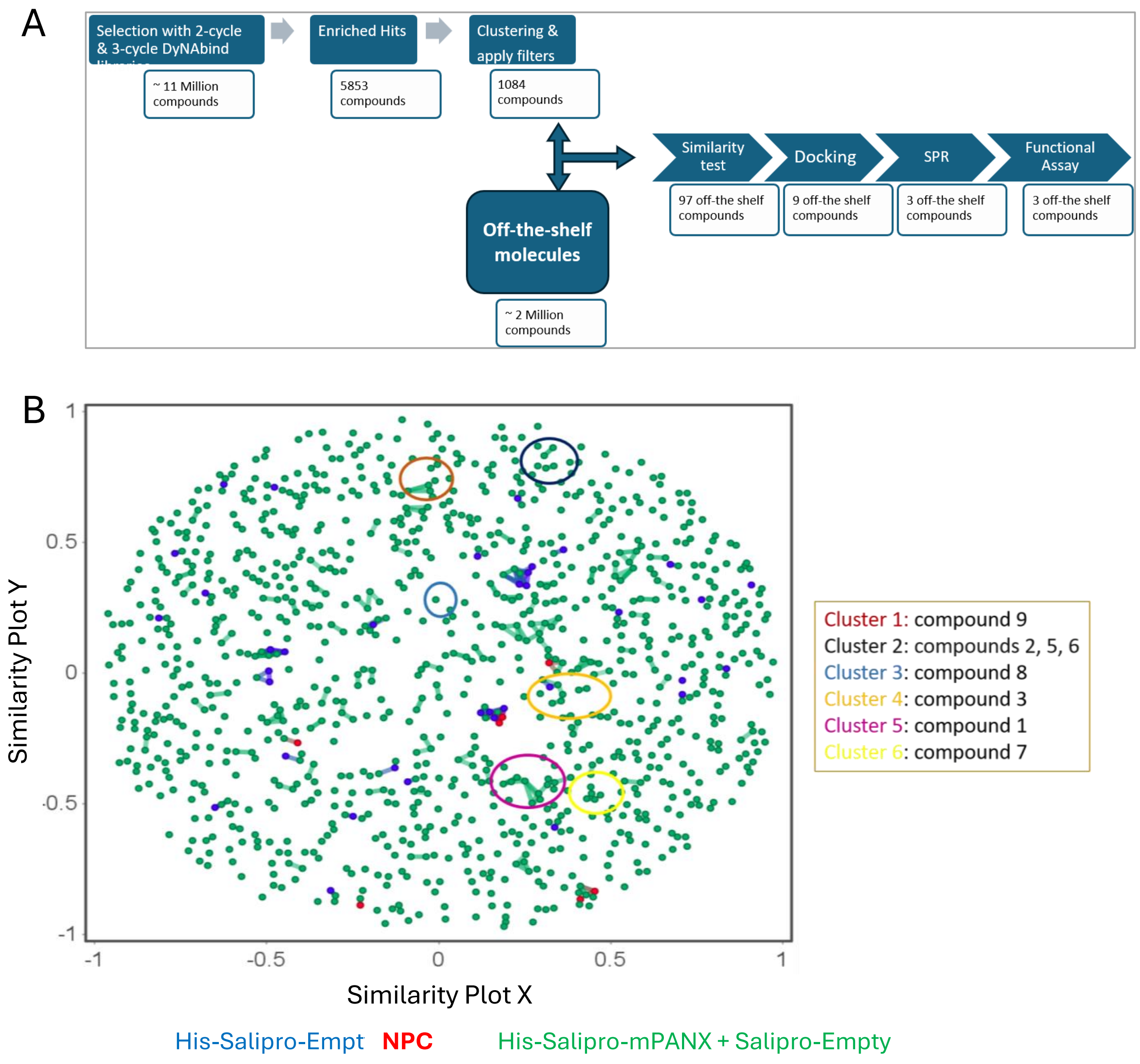


**Supplementary Figure 2.** DEL selection workflow. (A) Workflow for the prioritization of enriched hits for off-the-shelf validation. From the top 5,853 enriched compounds, clustering based on Morgan fingerprints (Tanimoto > 0.8) identified structurally related groups. After filtering out low-confidence hits, defined by low on-target enrichment, high off-target activity, or structural singletons, 1,084 candidate binders remained. Similarity screening against a 2-million-compound Mcule library (similarity > 0.85) yielded 97 off-the-shelf analogs. All 97 compounds were subsequently prioritized and docked to PANX1 protein structures. Nine compounds with favorable docking profiles were selected for purchase and subjected to SPR validation and functional assays. (B) Cluster analysis of the selected DEL compounds. Chemical similarity plots were used to visualize the most significantly enriched hits from the DEL selections. The position of each dot along the XV axis indicates chemical similarity. Each dot represents a detected hit compound showing high enrichment under one of three conditions: His-tagged pannexin immobilized on beads with 10× molar excess of untagged nanoparticles (His-Salipro-mPANX + Salipro-Empty, green), nanoparticles only (His-Salipro-Empty, blue), or negative control (NPC, red). Lines connecting dots indicate particularly strong structural similarity. Representatives from clusters highly enriched under the target condition were further analyzed via molecular docking and biophysical binding assays. Compounds 1–9 are commercially available and reflect the properties of the clusters represented by the colored circles.

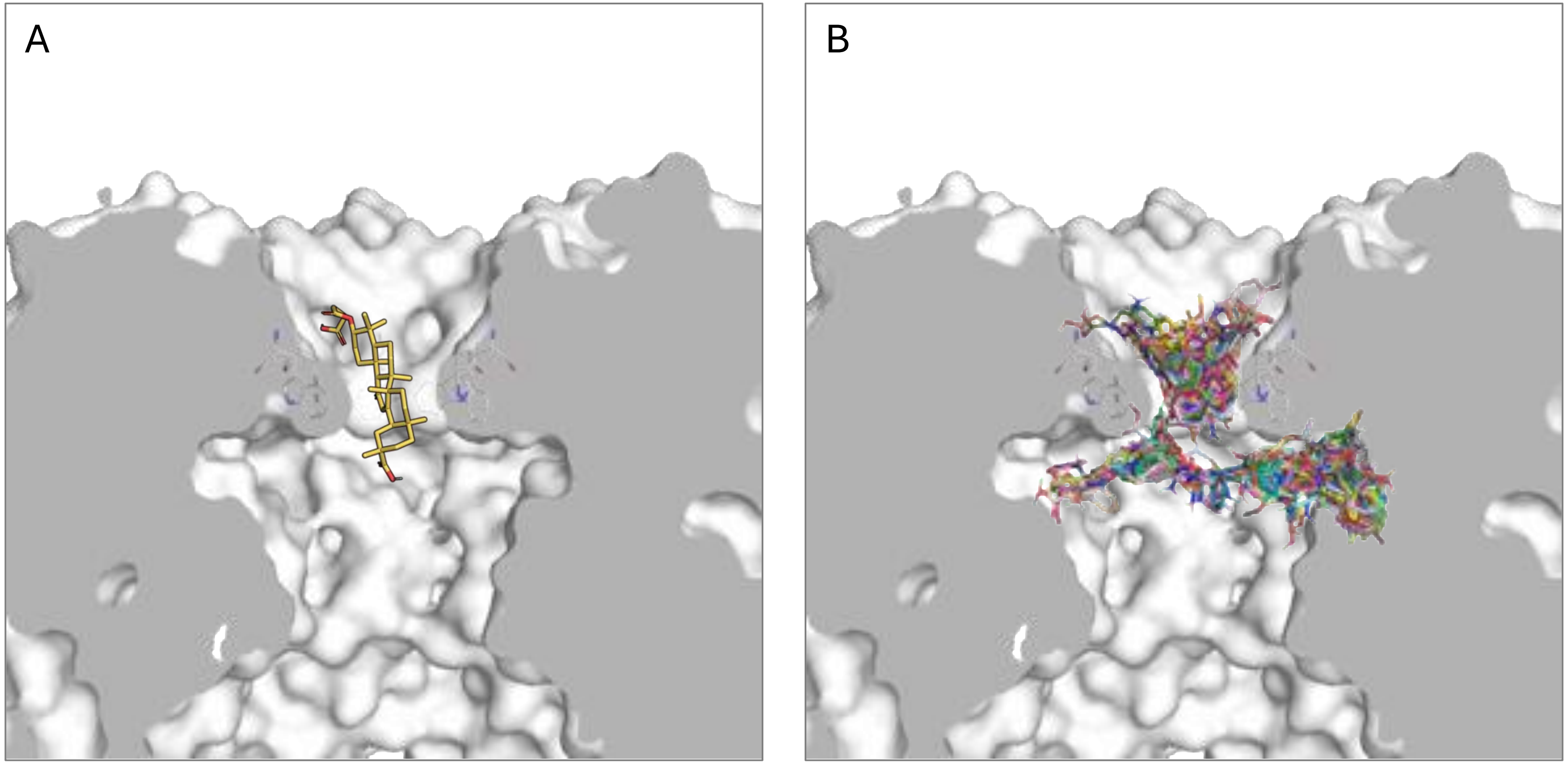


**Supplementary Figure 3.** Docking poses of the well-established PANX1 inhibitor carbenoxolone (A) and selected DEL-derived compounds (B) on mPANX1. The channel pore of mPANX1 is displayed as a surface cross section while the tryptophan belt and the docked compounds are displayed as sticks. All compounds were docked into a 30x30x30 $Å^3$ grid around the tryptophan belt's center of mPANX1 using AutoDock Vina v1.2.3 with the Vina forcefield and scoring function. Images were generated by PyMOL.

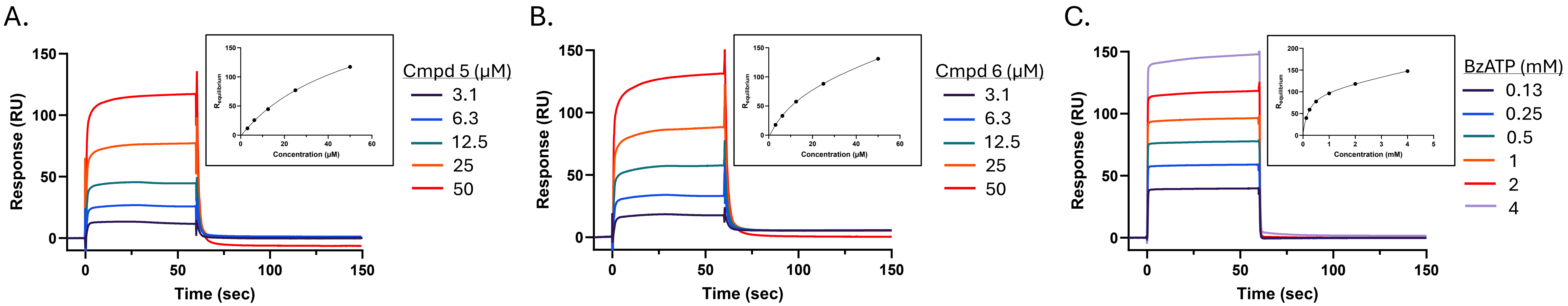


**Supplementary Figure 4.** Hit Compound Binding Profiles for hPANX1. Concentration series of (A) compound 5, (B) compound 6, and (C) benzoylbenzoyl-ATP were injected over a hPANX1 coated surface for 60 seconds followed by a 90 second dissociation time. Equilibrium analysis for each dataset is shown as an inset. Data is representative of n=2.

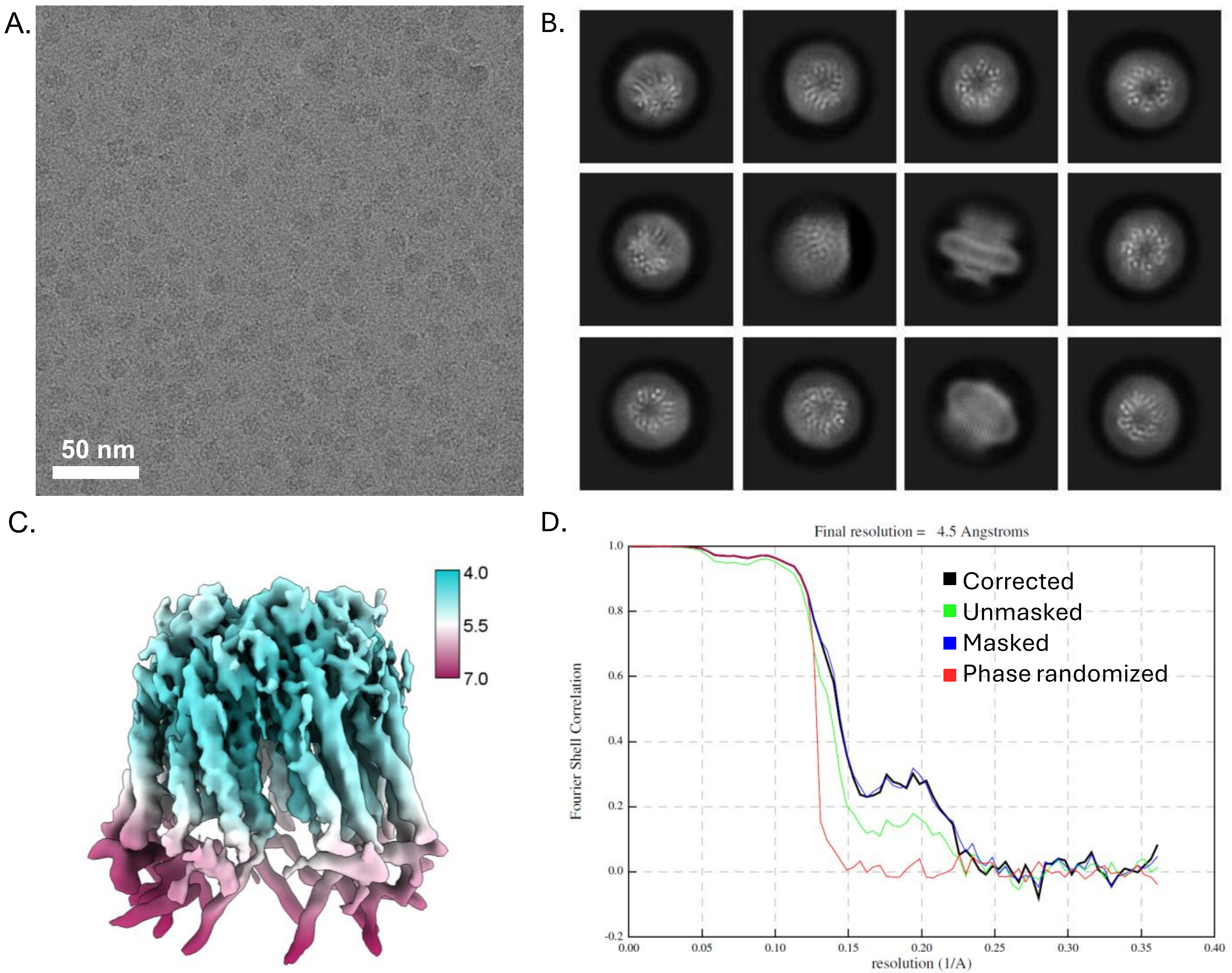


**Supplementary 5.** Cryo-EM refinement statistics and quality assessment. (A) Representative cryo-EM micrograph and (B) 2D class averages. (C) Cryo-EM map of Salipro-mPANX1 colored according to local resolution (in Ångström). (D) Fourier shell correlation (FSC) curve plot.

### Supplementary Table 1. Cryo-EM data collection and refinement statistics

| | EMDB: EMD-56448 |
|---|---|
| **Data collection and processing** | |
| Magnification | 165,000 |
| Voltage (kV) | 200 |
| Electron exposure (e–/Å$^2$) | 49.8 |
| Defocus range (µm) | -0.75 to -1.5 |
| Pixel size (Å) | 0.694 |
| Symmetry imposed | C1 |
| Initial particle images (no.) | 1,367,517 |
| Final particle images (no.) | 327,491 |
| Map resolution (Å) | 4.5 |
| FSC threshold | 0.143 |
| Map resolution range (Å) | 4.0-10.0 |
